# Integrative analysis of RNA, translation and protein levels reveals distinct regulatory variation across humans

**DOI:** 10.1101/018572

**Authors:** Can Cenik, Elif Sarinay Cenik, Gun W. Byeon, Fabian Grubert, Sophie I Candille, Damek Spacek, Bilal Alsallakh, Hagen Tilgner, Carlos L. Araya, Hua Tang, Emiliano Ricci, Michael P. Snyder

## Abstract

Elucidating the consequences of genetic differences between humans is essential for understanding phenotypic diversity and personalized medicine. Although variation in RNA levels, transcription factor binding and chromatin have been explored, little is known about global variation in translation and its genetic determinants. We used ribosome profiling, RNA sequencing, and mass spectrometry to perform an integrated analysis in lymphoblastoid cell lines from a diverse group of individuals. We find significant differences in RNA, translation, and protein levels suggesting diverse mechanisms of personalized gene expression control. Combined analysis of RNA expression and ribosome occupancy improves the identification of individual protein level differences. Finally, we identify genetic differences that specifically modulate ribosome occupancy - many of these differences lie close to start codons and upstream ORFs. Our results reveal a new level of gene expression variation among humans and indicate that genetic variants can cause changes in protein levels through effects on translation.

## Introduction

Deciphering the molecular mechanisms that underlie human variation is essential for understanding human diversity and personalized medicine. To date, genetic variants that affect protein function in humans have been well studied, but those that control protein levels are less well characterized. Yet, misregulation of protein levels can have profound consequences for human health. For example, transcriptional regulatory mutations that increase telomerase gene expression have been identified in ∼70% of melanoma patients (Horn et al., 2013; Huang et al., 2013) and are frequent in several other cancers (Huang et al., 2013). Similarly, changes in the protein levels of Shank3, neuroligins and neurexins have been linked to autism spectrum disorder, schizophrenia and learning disorders (Darnell, 2011). Therefore, understanding how RNA levels and translation efficiency control protein levels on an individual basis is required not only for understanding human phenotypic diversity, but also for personalized medicine as thousands of human genome sequences become available.

Protein expression is determined at many levels including (1) RNA expression, (2) translation efficiency and (3) protein stability. Recent studies have begun to unravel the extent of human variation at RNA levels and its control through transcription factor binding sites, and chromatin (Kasowski et al., 2010, 2013; McDaniell et al., 2010; Montgomery et al., 2010; Pai et al., 2012; Pickrell et al., 2010; Stranger et al., 2007; Westra et al., 2013). However, protein levels often correlate poorly with RNA expression (Ly et al., 2014; Vogel and Marcotte, 2012). Translation efficiency, i.e. the number of proteins synthesized per mRNA, has been suggested to account for a large component of the unexplained variation in protein levels (Marguerat et al., 2012; Schwanhäusser et al., 2011). While recent studies in yeast have begun to address the genetic control of translation efficiency (Albert et al., 2014; Artieri and Fraser, 2014; McManus et al., 2014; Muzzey et al., 2014), little is currently known about variation in translation efficiency and its genetic determinants in humans. Further, an integrated view of how expression is controlled at many different levels is lacking in humans.

Here, we utilized RNA-seq and ribosome profiling to identify ribosome occupancy profiles. Ribosome profiling involves RNAse digestion of unprotected RNA and isolation of ribosome-bound mRNA segments. The protected mRNA is sequenced to reveal the amount of ribosomes per message and deduce translation efficiency in conjunction with RNA-seq data. We integrated these measurements with quantitative proteomics to reveal a more comprehensive view of the variation in gene expression programs across a diverse set of humans.

## Results

### Measuring Ribosome Occupancy across Individuals at a Global Scale

To measure genome-wide ribosome occupancy of mRNAs, we first improved and adapted the ribosome profiling protocol (Ingolia et al., 2009, 2011, 2012) for lymphoblastoid cell lines (LCLs) (Figure 1A). A critical step in ribosome profiling method involves RNase digestion of unprotected RNAs before isolating and sequencing ribosome-associated mRNAs. While optimizing the protocol, we observed that RNAse I digestion caused extensive degradation of ribosome integrity in cytoplasmic lysates (Figure 1B-C, Supplemental Figure S1A). The loss of polyribosome signal was not accompanied by a corresponding increase in monosome signal (Figure 1C; Supplemental Figure S1A) but rather a shift towards lighter fractions indicative of free and degraded RNAs (Figure 1C; Supplemental Figure S1A). This suggests that the RNase I treatment is leading to a loss in ribosome integrity in addition to producing the expected 80S ribosome footprints (i.e. monosomes).

**Figure 1.**
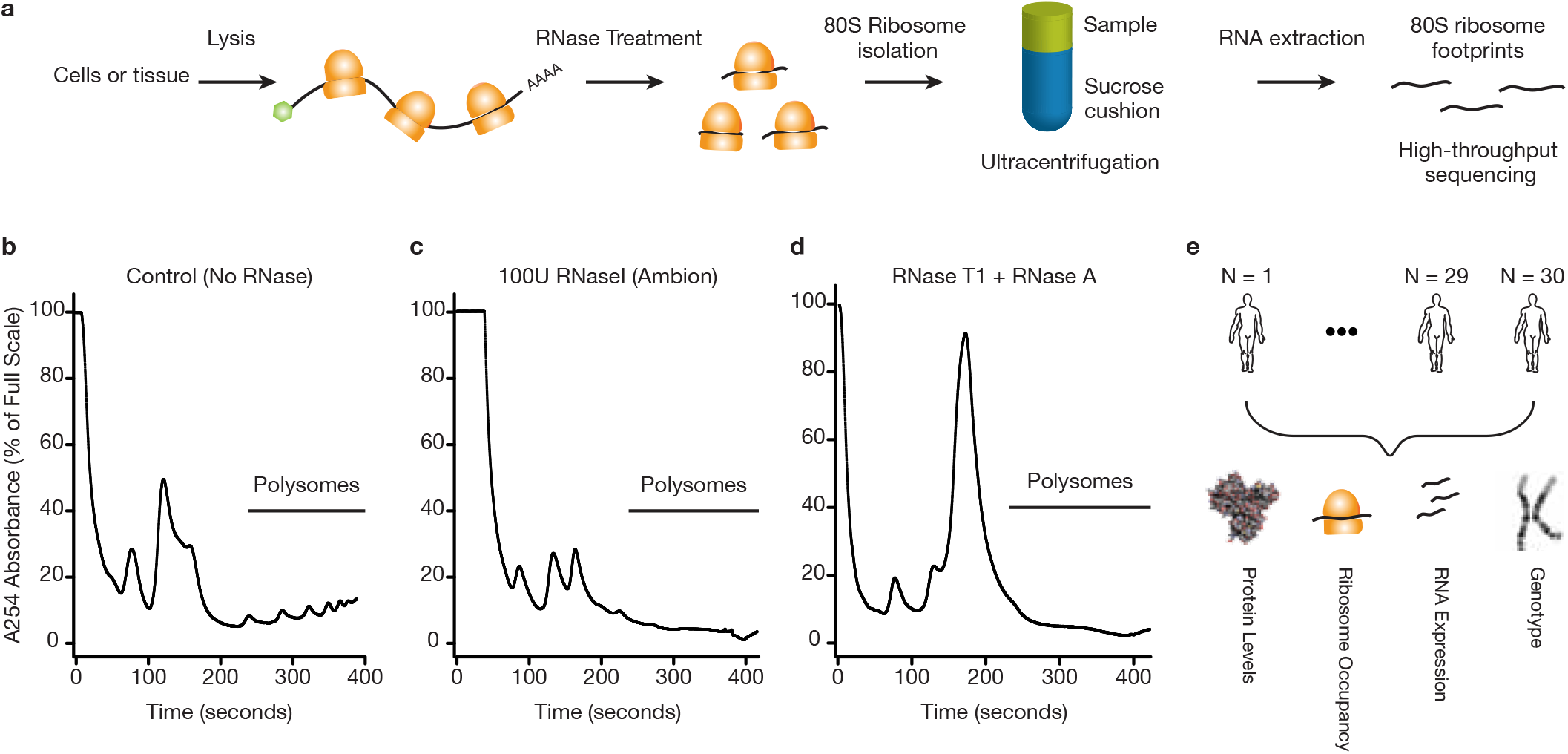
Choice of RNase is critical for generating ribosome profiling data. **(A)** A schematic representation of ribosome profiling strategy was shown. A key step is the digestion of unprotected RNA segments with an RNase. The ribosome protected RNA segments are isolated using a sucrose cushion and prepared for high throughput sequencing **(B)** Human lymphoblastoid cells (GM12878) were lysed in the presence of cycloheximide. The samples were ultracentrifuged through a 10-50% sucrose gradient. Samples were fractionated while continuously monitoring absorbance at 254nm. A representative polysome profile is shown **(C)** Samples were prepared for ultracentrifugation as in panel (b) with the following exception. The cleared lysate was incubated with 100 Units of RNase I (Ambion) for 30 minutes at RT before the ultracentrifugation. (**D)** Samples were prepared as in panel (c), except 300 Units of RNase T1 (Fermentas) and 500ng of RNase A (Ambion) was used for the RNase digestion step. A complete digestion of polysomes into monosomes was observed **(E)** Schematic representation of the datasets used in the current study. Genotype, ribosome profiling, RNA-Seq and mass spectrometry based proteomics data were collected from lymphoblastoid cells derived from a diverse group of 30 individuals.

We tested whether other RNases could alleviate this problem and found that treatment with RNase A and RNase T1 (which collectively cut after C, U and G) resulted in complete digestion of polyribosomes into monosomes (Figure 1D; Supplemental Figure S1A). Recent work in Drosophila and other systems also reported the importance of optimizing nuclease digestion to generate robust ribosome profiling data (Dunn et al., 2013; Ricci et al., 2014). Using our optimized ribosome profiling protocol, we generated ribosome occupancy maps for LCLs obtained from thirty individuals of diverse ethnic backgrounds: five Europeans, two Asian and twenty-three Yorubans with significant genetic diversity (Figure 1E). These lines were chosen because a) their genomes have been sequenced (1000 Genomes Project Consortium et al., 2012; International HapMap 3 Consortium et al., 2010), b) their relative protein and RNA levels have been previously measured (Khan et al., 2013; Wu et al., 2013), and c) they can be grown in large quantities. Importantly, the ribosome occupancy maps were based on at least two replicate samples for the majority of individuals. In parallel, we generated 44 deep RNA-Seq libraries (with a median of ∼12M uniquely transcriptome mapped reads) from the same cells and combined these with those from previous work (’t Hoen et al., 2013; Lappalainen et al., 2013; Pickrell et al., 2010) thereby providing multiple RNA-seq replicates for most individuals.

We leveraged replicate measurements to assess data quality and its dependence on several parameters including alignment strategy, mRNA enrichment method, PCR artifacts, gene length normalization and batch effects (Supplemental Figure S2A-E, see Methods). In addition to verifying the high quality of the data, replicate measurements also enabled modeling of gene-specific variance in RNA expression and ribosome occupancy per individual, allowing robust derivation of individual-specific translation efficiency estimates. Specifically, we developed a linear modeling based approach to regress out the effects of RNA expression from ribosome occupancy measurements to calculate translation efficiency (see Methods).

Finally, for 28 individuals studied here, we previously measured relative protein abundances via isobaric tag-based quantitative proteomics using the same cell lines (Wu et al., 2013). In total, we present a combined analysis of 133 high-throughput sequencing libraries (83 RNA-Seq and 50 ribosome profiling libraries) and extensive protein expression measurements (Figure 1E).

### Integrative Analysis of RNA Expression, Ribosome Occupancy and Protein Levels

We first considered the general relationship between RNA expression, ribosome occupancy, and protein expression. As expected, RNA expression and ribosome occupancy were highly correlated (Supplemental Figure S2F; Spearman *ρ*= 0.87, p-value < 2.2e-16; outlier robust correlation 0.88 using Donoho-Stahel estimator), albeit still lower than biological replicates of RNA expression data (Spearman *ρ* ∼ 0.98, p-value < 2.2e-16) indicating that control of ribosome occupancy levels is distinct from RNA levels. Importantly, ribosome occupancy correlated better with protein levels than RNA expression correlated with protein levels (Supplemental Figure S2F-G; Spearman *ρ* of 0.54 and 0.43, Donoho-Stahel estimator based correlation coefficient 0.56, and 0.42, respectively; permutation test for difference in correlation coefficient p-value < 10^-4^).

While the correlation analysis reveals pairwise relationships, the interdependencies between RNA expression, translational efficiency, and protein levels are not captured. For example, some genes with high protein levels can have low RNA expression but very high translation efficiency, yielding a decreased correspondence between RNA expression and protein levels. To reveal such interdependencies, we utilized self organizing maps (SOM), an integrative machine learning method that is robust to noise and allows assessment of all relationships simultaneously (Figure 2A) (Kohonen, 1990; Wehrens and Buydens, 2007). Since SOMs are sensitive to differences in mean and variance of the input variables, we first converted each measurement into their relative rank order expressed as percentiles ensuring equal weighting of the input variables for the SOM training (see Methods). After training, each neuron within the SOM contains genes that share a similar pattern of expression and protein level (Figure 2A; Supplemental Figure S2H).

**Figure 2.**
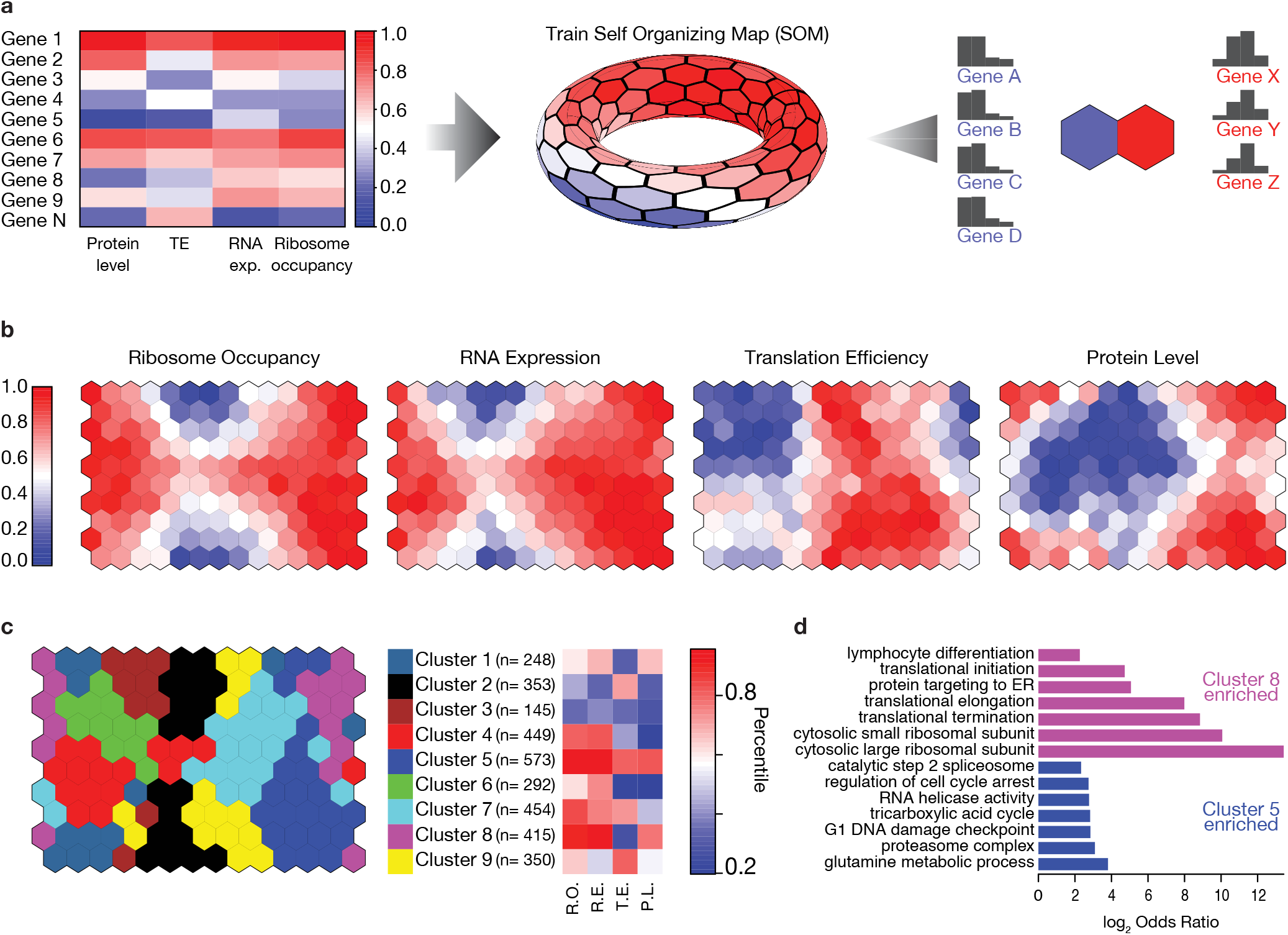
Ribosome occupancy correlates better with absolute protein levels than RNA expression and protein levels. **(A)** A self-organizing map (SOM) was trained using ribosome occupancy, RNA expression, translation efficiency and protein levels. These measurements were converted into their relative rank order before training. After training, each neuron in the SOM contains several genes sharing similar expression patterns. **(B)** Four different colorings of the trained SOM depict the mean ribosome occupancy, RNA expression, translation efficiency or protein levels for each neuron. **(C)** Neurons of the SOM were grouped using affinity propagation clustering (Frey and Dueck, 2007). Shared coloring between nodes indicates membership to the same cluster. For each cluster, the mean rank in ribosome occupancy (RO), RNA expression (RE), Translation efficiency (TE) and protein level (PL) was shown for the representative neuron of the cluster. The number of genes in each cluster (n) was shown. **(D)** For four out of nine clusters, significantly enriched gene ontology (GO) terms were identified (FuncAssociate (Berriz et al., 2009) permutation based corrected p-value < 0.05; Supplemental Table S1). For two clusters (5 and 8), selected GO categories were shown (log2 odds ratio; Supplemental Table S1 for the full list of enriched terms).

The emerging map recapitulated the general relationship between protein levels with RNA expression and ribosome occupancy as evidenced by the high correlations across neurons (Figure 2B). We further grouped neurons in the SOM using a clustering approach (affinity propagation clustering (Frey and Dueck, 2007). This approach uncovered nine clusters in the SOM, revealing the distinct relationships between RNA expression and translation efficiency in determining protein levels. For example, genes in Cluster 6 have relatively high RNA expression but do not reach high protein levels as they are translationally repressed (Figure 2C).

We then examined functional (GO term) enrichments (Berriz et al., 2003, 2009) across the different clusters within the SOM and found specific functional enrichments for four of the nine clusters (Figure 2C; Supplemental Table S1). Genes with high translation efficiency and high protein levels were enriched for diverse functional categories such as the proteasome complex, glycolysis, mRNA splicing, and DNA damage checkpoint (Supplemental Table S1; selected examples are shown in Figure 2D; p-value < 0.05 for all categories using permutation based multiple hypothesis correction). Conversely, genes associated with translation and cytosolic ribosome constituents were enriched among those that exhibited very high RNA and protein levels despite having lower translation efficiencies (Supplemental Table S1; Figure 2D; p-value < 0.05 for all categories using permutation based multiple hypothesis correction), raising the possibility that higher protein stability or feedback mechanisms on translation efficiency modulate the levels of translation components. These findings indicate that some sets of functionally coherent genes adopt alternative strategies to achieve their respective steady-state protein levels.

### Gene Expression Variability between Individuals

We next focused on how ribosome occupancy and RNA expression differ between individuals. We leveraged replicate measurements and identified genes with *significant* inter-individual variance in RNA expression or ribosome occupancy, exceeding technical noise. We found that ∼27% of genes had statistically significant inter-individual variation in RNA expression compared to only ∼7% of genes that had detectable variation in ribosome occupancy (Figure 3A; 3B; Holm’s method adjusted p-value < 0.05 based on a simulation based likelihood ratio test). Consequently, ∼20% of all genes exhibit inter-individual RNA expression variation which is not reflected in ribosome occupancy. These results were not explained by different sensitivities of the measurements (Supplemental Figure S3A). These results were also consistent when restricting the analysis to only the Yoruban individuals or when excluding RNA expression data not generated by our laboratory, indicating the robustness of the results.

**Figure 3.**
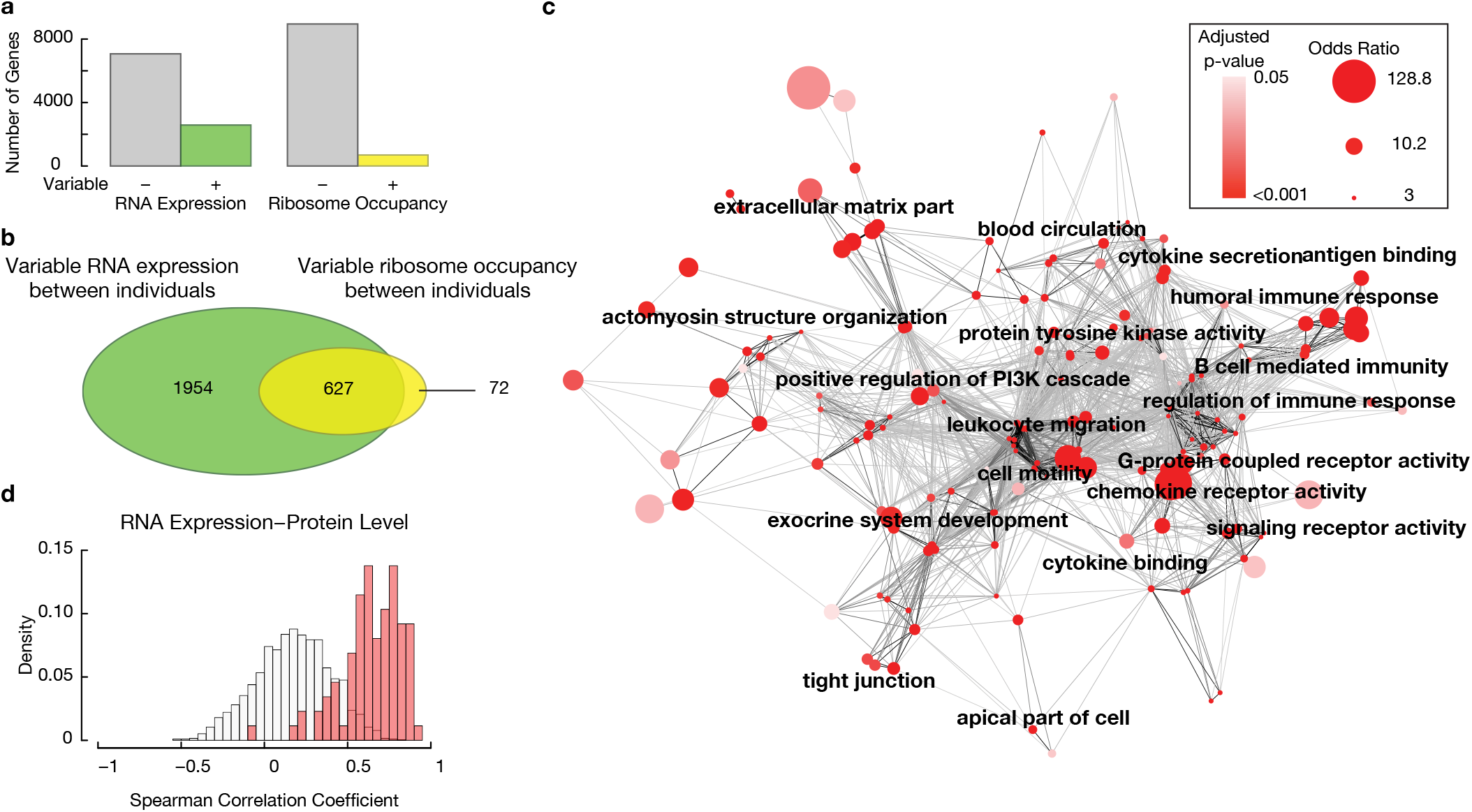
Identification of genes with significant inter-individual variability in RNA expression and ribosome occupancy improves the ability to identify personal differences in protein levels. **(A)** Ribosome occupancy and RNA expression was modeled using a linear mixed model treating individuals as a random effect and mean expression as the fixed effect. A simulation based exact likelihood ratio test (Scheipl et al., 2008) was used to compare the linear mixed model to a linear model that did not include the individual as a predictor. The number of genes that show significant inter-individual in RNA expression or inter-individual variation in ribosome occupancy was plotted (Holm’s corrected p-val < 0.05). **(B)** The Venn diagram depicts the overlap between the two groups **(C)** Enriched gene ontology (GO) terms among genes with significant inter-individual variation in both RNA expression and ribosome occupancy was determined using FuncAssociate (Berriz et al., 2009). Cytoscape (Smoot et al., 2011) was used to visualize the enriched GO terms (permutation test corrected p-value <0.05, odds ratio > 3; Supplemental Table S2). Nodes correspond to GO terms and are colored by the corrected p-value. The size of the node is proportional to the logarithm of the odds ratio. The similarity between GO terms was quantified using Kappa similarity. The strength of the similarity was visualized using darker edge colors (see Methods). An edge-weighted spring embedded layout was shown. **(D)** For each gene, Spearman correlation was calculated between individual specific RNA expression and relative protein levels. The distribution of the correlation coefficients was plotted as a density. Genes that showed significant variation in both RNA expression and ribosome occupancy between individuals were plotted with red bars and genes without detectable variation in RNA expression and ribosome occupancy were shown with white bars.

Genes that exhibited significant inter-individual variation in both RNA expression and ribosome occupancy were highly enriched for gene ontology terms including: “chemokine receptor activity”, “complement activation”, “ leukocyte migration”, “antigen binding” (Figure 3C; Supplemental Table S2; p < 0.05 permutation based multiple hypothesis correction), indicating a role in immune functions. Consistently, protein levels that exhibit the most variation between individuals were previously shown to be enriched for “immune system process” (Wu et al., 2013). These functional categories are highly specific to the function of the studied cell type, LCLs (Figure 3C; Supplemental Table S2). Given that genes with significant inter-individual variation were directly pertinent to the function of the cell line studied here, it is likely that carrying out similar studies in other cell types will expand the set of genes whose expression levels differ significantly between individuals.

Within genes that exhibited significant inter-individual variation in both RNA expression and ribosome occupancy, we identified three subsets. Within the first subset the variability in RNA expression was comparable to variability in ribosome occupancy (Supplemental Figure S3E). This first subset contained 54% of all genes exhibiting inter-individual variation in both RNA expression and ribosome occupancy. The second subset consisted of genes that had higher RNA level variability that is buffered at the ribosome occupancy level. This subset encompassed nearly twice as many genes as the third subset where ribosome occupancy variability was higher than that of RNA expression (Supplemental Figure S3E). These results were consistent with the observation that for many genes inter-individual RNA expression variation is not reflected in ribosome occupancy.

A small, but interesting fraction of genes (0.7%) exhibited differential ribosome occupancy between individuals with no apparent differences at the RNA-level (Supplemental Table S3). These were enriched in genes coding for proteins involved in “cellular response to chemical stimulus” and the “Golgi apparatus” (Supplemental Table S2; p < 0.05 permutation based multiple hypothesis correction). These results suggest that translational control may play important roles in cellular signaling, whereby rapid cellular responses are often required. Phosphatidylinositol 3-kinase subunit Gamma (*PIK3CG*), a promising target for drug development against inflammatory disease and cancer (Ghigo et al., 2010; Subramaniam et al., 2012), was among the genes with significant inter-individual variation in ribosome occupancy. Clinical Trials are currently investigating inhibition of *PIK3CG* in B-and T-cell leukemias and lymphomas (Clinical Trials NCT01871675; NCT02004522). Given that LCLs are derived from B-cells, stratifying patients on protein level variability of this gene may be important in drug development and dosage determination.

### Relationship between Individual Differences in Protein Levels, Ribosome Occupancy, and RNA Expression

An outstanding question in understanding phenotypic variation is how individual-specific protein levels relate to corresponding differences in gene expression. We previously measured relative protein levels for ∼6000 proteins using the same cell lines (Wu et al., 2013). As expected, the protein level measurements were skewed towards genes that are more highly expressed and translated (Supplemental Figure S3B, Wilcoxon rank sum test p < 2.2e-16). We first calculated the correlation between RNA expression and the corresponding protein level across individuals. Consistent with previous results (Wu et al., 2013), the median correlation coefficient was 0.22, with 20% of genes showing a statistically significant correlation (Figure 3D; Spearman correlation coefficient, 10% FDR using Benjamini-Hochberg method; Supplemental Figure S3C).

We next repeated this analysis for the set of genes that we identified as having significant RNA expression variability between individuals. Among this subset, relative protein levels and RNA expression had a median correlation coefficient of 0.43 (Spearman correlation coefficient; Supplemental Figure S3D), indicating a partial correlation between RNA and protein variability.

Finally, we tested whether joint measurement of RNA expression and ribosome occupancy improved this correspondence. Specifically, we considered genes that exhibit significant inter-individual variation in both ribosome occupancy and RNA expression. Strikingly, 93% of these genes had statistically significant correlation between differences in protein levels and RNA expression (Figure 3D; Spearman correlation coefficient, 10% FDR using Benjamini Hochberg method) with a median correlation coefficient of 0.67; and Supplemental Figure S3C). The large difference between the correlation coefficients indicates that measuring both ribosome occupancy and RNA levels simultaneously greatly improves the ability to identify gene expression variability between individuals that will eventually result in personal differences in protein levels.

### Genetic Determinants of Variability in Ribosome Occupancy

We next investigated whether genetic differences between individuals were associated with the observed variation in gene expression, specifically at the ribosome occupancy level. We used two complementary approaches. First, we used the 21 unrelated individuals from the Yoruban population and conducted a cis-quantitative trait loci (cis-QTL) mapping approach. Using the cis-QTL mapping strategy, we identified significant association between single nucleotide polymorphisms (SNPs) and ribosome occupancy for 67 genes (Supplemental Figure S4A-D). While 34 out of the 67 roQTLs were not associated with significant differences in RNA expression (nominal association p-value >0.05), this analysis cannot conclusively show that these roQTLs are not associated with RNA expression. Overall roQTLs had consistent effects on RNA expression and protein levels (Supplemental Figure S4A-D; Spearman *ρ*=0.86, p < 2.2e-16). Consistent with recent work comparing two yeast strains (Albert et al., 2012), these results suggest that genetic effects that were propagated through RNA expression to ribosome occupancy caused consistent changes in protein levels for this set of genes.

### The Role of uORFs in Modulating Ribosome Occupancy

We next adopted a targeted approach that was both better powered and enabled detection of combined effects of multiple genetic variants on ribosome occupancy. We first analyzed variants modifying upstream open reading frames (uORFs), which can alter protein expression by regulating translation (Wethmar et al., 2014). In humans, approximately half of annotated transcripts contain uORFs, and presence of uORFs is widely polymorphic across individuals (Barbosa et al., 2013; Calvo et al., 2009; Waern and Snyder, 2013).

We correlated genetic alterations to uORF presence with ribosome occupancy of the main coding region and found 33 transcripts with significant association (Figure 4A-D, Supplemental Figure S4E-F; 5% FDR). One particular advantage of this targeted approach was the ability to detect changes to uORFs caused by multiple genetic variants. For example, two different SNPs in the ZNF215 gene result in merging of two uORFs by removing a stop codon (Figure 4D). Merging of the uORFs significantly increased ribosome occupancy of the main coding region (Figure 4D; p<0.001).

**Figure 4.**
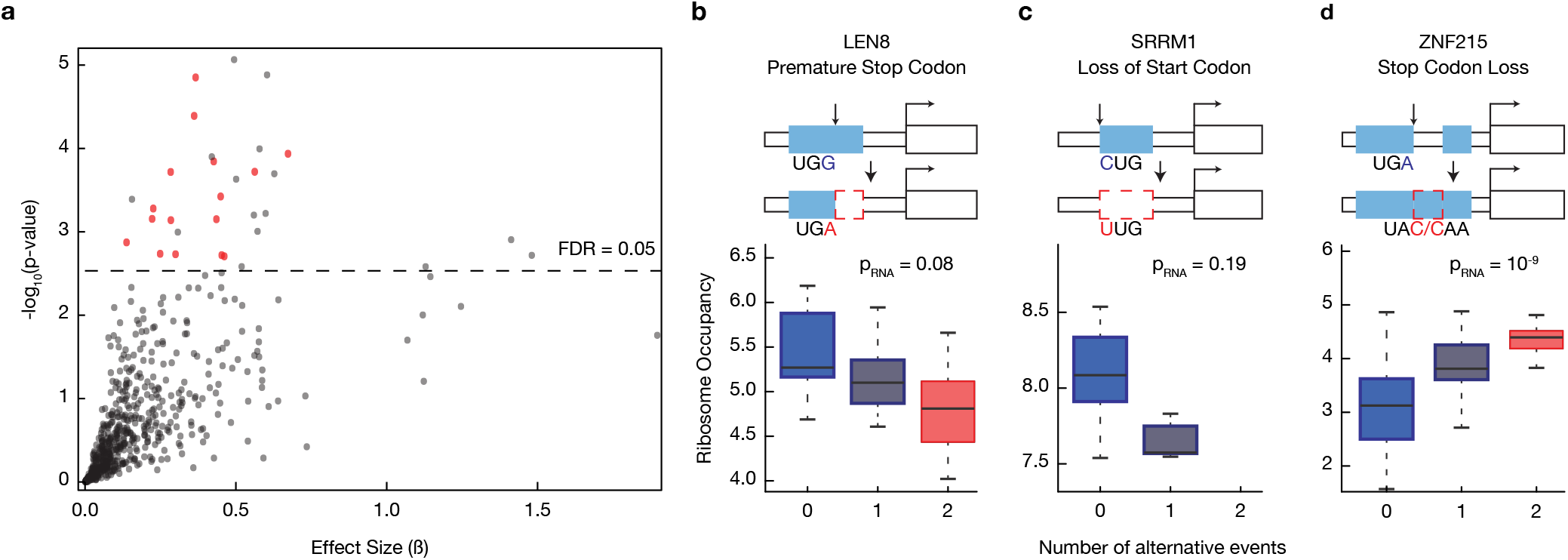
**Figure 4. Nucleotide variants that modify upstream ORFs can alter ribosome occupancy of the main coding region** **(A)** We identified single nucleotide polymorphisms that generate, delete or otherwise modify an upstream open reading frame (uORF). We tested whether changes to uORFs affected ribosome occupancy of the main coding region using a linear regression framework. The absolute value of the effect size from the regression was plotted against the p-value of association. For 17 uORF changes shown with red circles, the association was solely with ribosome occupancy (nominal p-value > 0.05 or opposite signed regression coefficients for RNA expression, and Supplemental Table S4 for robustness to population stratification and linear mixed model). **(B)** A SNP in the 5’UTR of LENG8 gene introduces a premature in-frame stop codon that shortens an existing uORF. This event results in lower ribosome occupancy of the main coding region, as shown in the boxplot (p_Ribo_ = 0.002). The horizontal bar reflects the median of the distribution and the box depicts the inter-quartile range. The whiskers are drawn to 1.5 times the inter-quartile range. **(C)** In another example, SRRM1, a SNP completely eliminates an existing uORF by removing its start codon. The loss of this uORF is associated with reduced ribosome occupancy of the main coding region (p_Ribo_ = 0.0004; p_RNA_= 0.19). **(D)** The reference sequence of ZNF215 gene has two short uORFs. Two different genetic variants eliminate the stop codon of the first uORF (UGA to UAC or UGA to CAA) resulting in merging of the two short uORFs into a single long uORF. The merging of the uORF significantly modulates both ribosome occupancy and RNA expression (p_Ribo_ = 0.0001; and p_RNA_ = 10^-9^ respectively).

In addition to impacting translational efficiency, nucleotide variants changing uORFs may alter RNA abundance. For example, they may change transcript stability by triggering nonsense-mediated decay (Kervestin and Jacobson, 2012). Alternatively, the variant or variants in linkage disequilibrium may alter transcriptional output as a proximal element downstream of the transcription start site. Of the 33 significant associations between changes in uORFs and ribosome occupancy, ∼52% (17 out of 33) also had a significant effect on RNA levels (nominal p-value < 0.05), indicating the presence of uORFs and RNA levels are often coupled. However, for 16 other genes the observed effect was solely on ribosome occupancy suggesting direct modulation of the translation efficiency of the main reading frame (Figure 4A-B, Supplemental Table S4). We further observed that presence of uORFs could be associated with both increased and decreased ribosome occupancy of the main coding region (Supplemental Figure S4G). We verified the robustness of these results by limiting the analysis to data from Yoruban individuals and employing an alternative statistical framework based on linear mixed models (Supplemental Figure S4E-F and Supplemental Table S4). These results reveal that natural genetic variation within the human population can specifically cause personal differences in translation through changes to uORFs.

### The Role of Sequences Surrounding the Start Codon in Modulating Ribosome Occupancy

We next analyzed the Kozak sequence, the region surrounding the start codon for its effect on translation efficiency (Figure 5A). Previous work has suggested that Kozak sequence is important for both start codon selection and translation efficiency of specific transcripts (Kozak, 1987). However, the extent and impact of natural genetic variation affecting the Kozak sequence and the global effect of the Kozak region on translation efficiency have not been studied.

**Figure 5.**
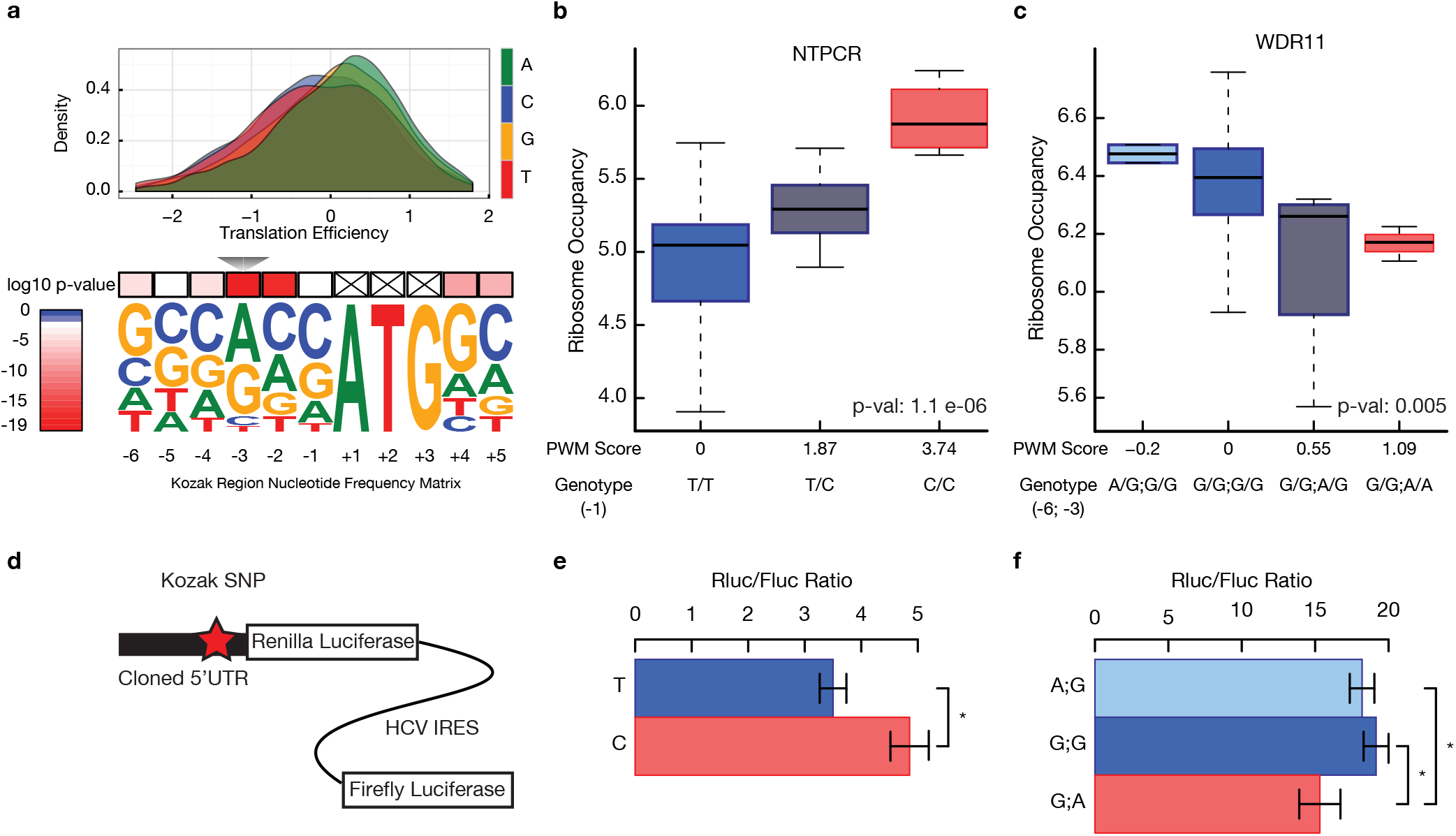
Nucleotide variants modulating the sequence around the translation initiation site alter translation efficiency. **(A)** Kozak region is defined as the six nucleotides preceding and two nucleotides following the start codon. The derived position weight matrix was visualized using WebLogo (Crooks et al., 2004). The upper panel shows the effects of each nucleotide at the -3 position on translation efficiency. The effect of nucleotides on translation efficiency was tested using Kruskal-Wallis test. **(B)** The effect of a Kozak region variant on the ribosome occupancy of NTPCR was assessed using a linear model (p-value = 1.1e-6). A boxplot was used to visualize the distribution of ribosome occupancy for individuals with given genotypes. The horizontal bar reflects the median of the distribution and the box is drawn to depict the inter-quartile range. **(C)** WDR11 had two naturally occurring SNPs in its Kozak region. An additive model was adopted to calculate the change in the position weight matrix score of the Kozak region. **(D)** 5’UTRs with or without Kozak variants were cloned into a translation efficiency reporter. The reporter expresses a biscistronic mRNA where the Renilla luciferase is translated under the control of the cloned 5’UTR and the Firefly luciferase is translated under the control of Hepatitis C virus (HCV) internal ribosome entry site (IRES). **(E)** and **(F)** The ratio of Renilla to Firefly luciferase activity was plotted for NTPCR (e) and WDR11 (f). Error bars represent s.e.m. The difference between the ratios was assessed using a two sided two sample t-test (* denotes p-value < 0.05).

We first determined whether certain positions of the Kozak sequence have a global effect on translation efficiency. We found a highly significant and large effect of the nucleotides at position -3 and at position -2 on translation efficiency (Figure 5A; Supplemental Figure S5A; Bonferroni adjusted Kruskal-Wallis test p=5.7e-20 for position -3; p=1.2e-17 at position -2). Additionally, the two nucleotides immediately after the start codon had statistically significant effects on translation efficiency (Supplemental Figure S5A; Bonferroni adjusted p < 2.8e-7). While previous work using reporter systems anticipated the significance of these features (for example, Kozak, 1987), our analyses highlight the role of sequence composition near the ATG in modulating translation efficiency of endogenous genes at a global scale.

The extent and potential role of natural variation that might alter the Kozak sequence across the genome remains largely unexplored in the human population (Xu et al., 2010). Among the set of individuals studied here, there were ∼150 genetic variants altering the Kozak region in at least three individuals. ∼65% of Kozak region variants reduced the PWM score of the reference sequence (Supplemental Figure S5B). This effect was even more pronounced for Kozak variants that were observed in a single individual. 77% of these reduced the PWM score of the reference sequence suggesting that selective pressure may be acting to optimize the Kozak sequence.

We next tested the effect of these variants on ribosome occupancy of the main coding region. We utilized the position weight matrix for the Kozak region to score the impact of each variant on the Kozak strength (Figure 5A). We found nine genes with Kozak variants that modified ribosome occupancy significantly with no significant effect on the RNA levels (Figure 5B-C; 10% FDR using Benjamini-Hochberg correction; RNA expression association p-value > 0.01; Supplemental Figure S5C; using a conservative linear mixed model, two of these genes had p < 0.01), indicating the presence of variants specifically affecting translation efficiency.

Finally, to directly examine the role of genetic variation on translation efficiency, we used reporter assays (Jang et al., 1988) for six genes. These included four genes with Kozak region variants and two genes with uORF variants. We cloned the reference 5’UTR or the variant 5’UTR with a single base change at the Kozak region or the uORF upstream of a Renilla luciferase and transfected the resulting constructs into human HEK 293 cells. To normalize differences in RNA expression and transfection efficiency, an HCV internal ribosome entry site driven Firefly luciferase was cloned after the Renilla stop codon and the ratio of the Renilla to Firefly luciferase was quantified. Differences in this ratio between the reference and variant 5’UTR-containing reporters for four genes recapitulated the results from our ribosome profiling data i.e. sequences that were associated with reduced translational efficiency also gave low luciferase ratios. These results provide an independent validation of our conclusion that natural genetic variation can modify sequences surrounding the start codon leading to personal differences in translation (Figure 5D-F; Supplemental Figure S5D).

## Discussion

This study demonstrates for the first time that translation efficiency varies among different individuals and that the nucleotides important for regulating translation efficiency can be mapped and identified. In several cases we uncovered the mechanisms controlling translation efficiency variation in humans. These included uORFs and sequences near the translation initiation sites. Other natural genetic variants were also identified but how they affect ribosome occupancy is currently not known. Our study revealed that genetic differences between individuals could lead to gene expression differences at the level of translation.

Most of our analyses were predicated on careful optimization of the ribosome profiling for LCLs and replicate measurements. In particular, our results reveal that RNase I is suboptimal for generating ribosome profiling data in certain mammalian cell including LCLs consistent with observations in several other systems (Figure 1 and Figure 1-figue supplement 1; Dunn et al., 2013; Ricci et al., 2014). Consequently, studies that do not specifically optimize ribosome profiling conditions may lead to misleading conclusions.

We leveraged replicate measurements to identify genes with significant variability in RNA expression or ribosome occupancy between individuals. We found that genes that exhibit significant variability in both RNA expression and ribosome occupancy were highly enriched for functions directly pertinent to LCLs such as immune response and leukocyte migration (Figure 3). Hence, extending this analysis framework to additional cell types or tissues will likely uncover more genes with variable expression between individuals.

We also found that many genes exhibited significant variability in RNA expression between individuals, yet this variability was not reflected at the translation level. While future studies will be needed to establish the generalizability of these results, recent yeast studies comparing translation level differences between two closely related yeast species (Artieri and Fraser, 2014; McManus et al., 2014) have similarly found that translation level differences compensated for RNA expression variability.

We then investigated the relationship between protein levels differences and variability in RNA expression and translation. We found that joint analysis of RNA expression and translation improved our ability to identify the extent of gene expression variation that will be reflected in protein levels, indicating a tight coupling of translation efficiency and protein levels. These analyses were skewed towards genes with higher expression levels due to missing protein level measurements (Supplemental Figure S3B). Hence, further improvements in proteomics technology will be needed to test the generalizability of our results to lowly expressed proteins. Despite the significant improvements obtained by joint analysis of ribosome occupancy and RNA level measurements, there remains unexplained variability in protein levels. One potential contributor to this discrepancy is variability in protein degradation rates (Vogel and Marcotte, 2012).

Our results demonstrate genes that have individual variability only in RNA expression are less likely to have corresponding differences at the protein level. Among this subset of genes, only 40% had statistically significant co-variation between RNA levels and protein levels (5% FDR). An important implication of this result concerns ongoing large-scale efforts that aim to identify genetic determinants of RNA expression (Battle et al., 2014; Lappalainen et al., 2013). These studies are in part motivated by the finding that most disease risk factors identified by genome-wide association studies lie in noncoding regions (Edwards et al., 2013). By linking genetic differences to RNA expression, these studies hope to uncover functional connections to disease states. Yet, our analyses suggest that the functional impact of RNA-level differences needs to be carefully considered to establish causal relationships to phenotype.

Recent consortium efforts measured RNA expression and genotype in large samples (∼900 in Battle et al., 2014; ∼500 samples in Lappalainen et al., 2013) leading to identification of trans-acting genetic effects on RNA expression. Interestingly, 85% of the trans-effects on RNA expression were found to be mediated by the effects of the associated SNP on a nearby gene (Battle et al., 2014), indicating that changes in regulators of RNA expression lead to differences in RNA levels of distant transcripts. Similarly, genetic variation in translation regulators are likely to have trans-effects on ribosome occupancy of many transcripts. For example, levels of global regulators of translation such as mTOR, and translation initiation factors (e.g., eIF4E) have the potential to modulate the translation of a large number of targets (Mamane et al., 2007; Thoreen et al., 2012). In fact, a recent study that compared translation in two different strains of yeast suggested that the relative importance of trans-effects on translation is comparable to that for RNA levels (Albert et al., 2014). We believe that our study will pave the way for future efforts that will explore translation variability with similarly large sample sizes. Such future studies will likely uncover trans-acting and additional cis-acting genetic variants associated with translation, and reveal the contribution of population-level variation to translation variability.

While our manuscript was in preparation, a recent study (Battle et al. 2015) reported associations between RNA expression, ribosome occupancy and protein. However, we note critical limitations in the ribosome profiling data reported. As demonstrated in Figure 1 and recently by Miettinen and Björklund (2015), the nuclease digestion conditions employed in Battle et al. lead to severe ribosomal degradation and significantly lower monosome purity in ribosome profiling libraries. Moreover, the Battle et al. study design lacks replicate experiments, precluding both proper assessment of the reproducibility of their ribosome profiling experiments and derivation of robust translation efficiencies. Strikingly, the rank correlation between re-sequencing experiments was lower than 0.9 (Spearman *ρ*) for the majority of their ribosome profiling libraries (Figure S2A in Battle et al. 2015). In stark contrast, we consistently achieved >0.98 rank correlations between biological replicates using independently grown cells and independently prepared and sequenced ribosome profiling libraries. When comparing each ribosome profiling data to others in the study, 12.5% of Battle et al. samples had median rank correlation less than 0.85 (lowest 0.72) suggesting the presence of a significant number of low quality ribosome profiling samples. Conversely, all of our datasets had median rank correlations greater than 0.85 (lowest 0.89). Taken together, by leveraging higher quality ribosome profiling datasets and independent validation experiments we were able to identify genetic variants associated with ribosome occupancy but not RNA expression that were missed by Battle et al.

Our study revealed several genetic variants that control translation efficiency variation in humans, including those affecting the Kozak region and upstream open reading frames (uORFs). A particularly interesting question is the molecular mechanisms of these sequence-function relationships. An intriguing feature of genetic variants modifying uORFs on translation was the observation that both gain and loss events could lead to increased translation of the downstream open reading frame (Supplemental Figure S4G) consistent with previous work that implicated both positive and negative regulation of translation efficiency by uORFs (Brar et al., 2012; Waern and Snyder, 2013). Whereas several mechanisms have been implicated in negative regulation of translation efficiency by uORFs, including nonsense mediated decay (Kervestin and Jacobson, 2012), less is known about the mechanisms of positive regulation by uORFs. Recent work identified DENR-MCT1, a protein that catalyzes translation reinitiation downstream of certain uORFs (Schleich et al., 2014). It is possible that DENR-MCT1 or similar factors may act on subsets of uORFs to increase reinitiation frequency of the downstream ORF leading to higher translation efficiency.

Recent structural analysis of the yeast 48S translation initiation complex permitted an unprecedented view of the molecular environment of the start codon in eukaryotes (Hussain et al., 2014) revealing a potential mechanism by which Kozak region variants affect translation efficiency. Remarkably, this structural analysis revealed that eIF2a makes direct contact with the nucleotides at positions -2 and -3, the same two positions that our global analysis of Kozak variants highlighted as being the most important for translation efficiency (Figure 5A). Thus, our results provide functional evidence that these residues are of general importance for translational efficiency.

Together, these results demonstrate that genetic alterations in the human population and disease-associated mutations may penetrate to phenotype through changes in translation. In the era of personal genome sequencing, this information is crucial for understanding the role of genetic variants on gene expression, phenotypic traits and human disease susceptibility.

## Methods

A more detailed description of the experimental and computational methods are provided in Supplemental Materials.

### Cell culture and lysis

Human lymphoblastoid cell lines (LCLs) were obtained from Coriell Cell Repository and grown in 15% fetal bovine serum and 1% Pen-Strep. For replicate experiments, cells were grown separately to a density of 0.8-1.0 × 10^6 cells/mL. Approximately 10 million cells were pelleted at 250g at 4C and washed with PBS. The pellets were frozen in liquid nitrogen prior to cell lysis. Cells were lysed in 150ul of lysis buffer (20 mM Tris-HCl pH 7.5; 150 mM NaCl; 5mM MgCl2; 1mM DTT; 100 μg.ml^-1^ Cycloheximide; 1% Triton X-100; 25U.ml^-1^ Turbo DNase I).

## Sucrose gradient fractionation and ribosome profiling

For sucrose gradient fractionation of LCLs, 7 A260 Units of the cleared cell lysate were incubated with 100U or 500U of RNase I (Ambion) or 5U or 15U of RNase I (Epicentre) or 300 Units of RNase T1 (Fermentas) and 500ng of RNaseA (Ambion) for 30 minutes at room temperature. RNase digestion was stopped with 20 mM Ribonucleoside Vanadyl Complex (NEB: S1402S). No RNase control included 20 mM Ribonucleoside Vanadyl Complex in cell lysis buffer to inhibit any endogenous RNase activity. Samples were loaded on top of a 10-50% (Weight/Volume) sucrose gradient (20 mM Tris-HCl pH 7.5; 150 mM NaCl; 5 mM MgCl2; 1 mM DTT; 100 μg.ml^-1^ Cycloheximide) and centrifuged in a SW-41 rotor at 38,000 rpm for 2h30min at 4°C.

For ribosome footprint library preparation using LCLs, 7 A260 Units of the cleared cell lysates were incubated with 300U Units of RNase T1 (Fermentas) and 500ng of RNase A (Ambion). Instead of a sucrose gradient fractionation, a 34% (Weight/Volume) sucrose cushion was used. Sequencing library preparation was done as previously described with some modifications (Ingolia et al., 2012; see Extended Experimental Procedures).

## RNA-Seq experiments

Lymphoblastoid cell lines (LCLs) were grown at a density between 3*10^5-6*10^5 cells/ml. Total RNA was extracted using Trizol reagent according to the manufacturer’s instructions (Lifetechnologies), then purified using the Qiagen RNeasy kit (Qiagen, Valencia, CA) and treated with RNAse-free DNase (Qiagen, Valencia, CA). RNA integrity was checked with a Bioanalyzer (Agilent, Santa Clara CA) and only samples with an RNA integrity number (RIN) of > 9.5 were subsequently subjected to either ribosomal depletion or poly-A-selection. For ribosomal RNA depletion, 5 μg of purified total RNA was depleted of rRNAs using the Ribo-Zero Magnetic Gold Kit (Human/Mouse/Rat) (Epicentre Biotechnologies, Madison, WI) according to the manufacturer’s instructions. For poly-A selection, 10 μg of purified total RNA were enriched by performing two cycles of selection using the Dynabeads mRNA Purification Kit (Life Technologies). Stranded libraries were prepared following the dUTP protocol (Parkhomchuk et al., 2009). For each cell line, we generated 2 × 101bp paired end RNA-Seq data using two biological replicates of ribosomal RNA depleted and three biological replicates of poly-A-selected RNA.

## Sequence alignment and processing

To enable comparable analysis of different high throughput sequencing datasets, we employed a uniform alignment and preprocessing pipeline starting from raw reads in FASTQ format. Reads were aligned using Bowtie2 v.2.0.5 with a sequential strategy (Langmead and Salzberg, 2012). Human rRNA and tRNA sequences were downloaded from UCSC Genome Browser (hg19) repeatmasker track (http://www.repeatmasker.org). All reads mapping to these sequences were filtered out. The remaining reads were aligned to APPRIS principal transcripts (release 12) (Rodriguez et al., 2013) from the GENCODE mRNA annotation v.15 (Harrow et al., 2012). For all transcript level analyses, reads that map only to coding regions were used.

## Ribosome profiling sample identity verification

The cell line identity for all ribosomal profiling libraries were verified by comparing empirically generated genotype calls to the reference genotypes. Specifically, we utilized samtools mpileup utility in combination with bcftools (Li et al., 2009) to generate genotype calls from the ribosomal profiling read alignments. Finally, a custom perl script was used to compare the number of perfect matches between empirically called genotypes and the reference genotype that was available from HapMap and 1K Genomes (see below).

## Genotype data and processing

Genome sequences were obtained from the 1000 Genomes Project pilot 2 trios and Phase1v3 (1000 Genomes Project Consortium et al., 2012; International HapMap 3 Consortium et al., 2010) for 27 of the 30 individuals. The genome sequences of three YRI LCLs (NA19139, NA19193, and NA19201) were not available from the 1000 Genomes Project. For these three cell lines, genotypes were imputed from HapMap release 28 data data (International HapMap 3 Consortium et al., 2010; International HapMap Consortium et al., 2007) to the 1000 Genomes Phase1v3 reference panel (1000 Genomes Project Consortium et al., 2012).

## Processing variant calls for downstream analyses

We included all variant calls provided by both release and pilot datasets without additional score or source filtering. We subsetted all single nucleotide polymorphisms (SNPs) that overlap APPRIS transcripts, same set used in sequence read alignments as described above. We converted genome coordinates to 0-based transcriptome coordinates and retained all phasing information from the VCF files. About ∼8% of the variants in the pilot dataset were unphased, and for these variants, we randomly assigned the phase.

## Sequence data normalization and quality control

After accounting for PCR bias, and mRNA enrichment method for RNA-Seq, ∼9600 transcripts had a read count per million reads mapped (cpm) (as implemented in the edgeR package (McCarthy et al., 2012)) greater than one in at least 40 RNA-Seq libraries and 36 ribosome profiling libraries. These transcripts were retained for further analysis. We accounted for different library sizes by calculating normalization factors using trimmed mean of M values (Robinson and Oshlack, 2010), and estimated the mean to variance relationship in the data using the voom method (Law et al., 2014). We explicitly specified the individual identifier to indicate which libraries were replicates from the same individual while applying the voom method. The inverse variance weights obtained from the voom method were used in all analyses where applicable.

## Calculation of translation efficiency

When combined with RNA expression measurements, ribosome profiling enables the estimation of translation efficiency by capturing a snapshot of the transcriptome-wide ribosome occupancy. We treated ribosome profiling and RNA-Seq as two experimental manipulations of the RNA pool of the cell. By using the voom method, we explicitly modeled the mean to variance relationship and utilized inverse variance weights for the subsequent steps. This preprocessing strategy enabled the application of the well-established statistical methods in the limma R package (Smyth, 2005). Specifically, we used a linear model where the normalized expression values are dependent on the treatment (RNA-Seq or Ribosome profiling) and the individual identifiers.

## Self-Organizing maps for integrative gene expression analysis

We used SOMs to explore the relationship between protein levels and the three expression level measurements: RNA levels, ribosome occupancy and translation efficiency. SOMs rely on a suitable measure of distance between the transcripts for the clustering. To avoid skewing distance calculation due to difference in scale and variance of the expression measurements, expression levels and protein amounts were converted to percentiles using the empirical cumulative distribution function for each level. The kohonen R package (Wehrens and Buydens, 2007) was used for training the SOM with custom modifications to the plotting functions following (Xie et al., 2013). We then clustered the codebook vectors of the 140 units in the SOM using affinity propagation clustering (Frey and Dueck, 2007) as implemented in the apcluster R package (Bodenhofer et al., 2011).

## Gene set enrichment analysis

FuncAssociate 2.0 was used for gene set enrichment analyses (Berriz et al., 2009). The ensembl gene identifiers were used for the enrichment tests. The background gene list was explicitly defined as the set of all genes that could potentially be included in the query set. We defined significant enrichments as GO terms with an odds ratio greater than 2 and adjusted p-value < 0.05. P-value adjustment was carried out using a permutation method to account for the overlap between the GO terms. We calculated the Kappa Similarity Score between all pairs of significantly enriched GO terms. We retained edges between all pairwise GO terms whose Kappa similarity score was greater than 0.1. Enriched GO terms were visualized with Cytoscape (Smoot et al., 2011) using the edge-weighted spring embedded layout.

## Analysis of between individual variation in RNA expression and ribosome occupancy

Replicate measurements for RNA-Seq and ribosome profiling were used to determine inter-individual variance while controlling for platform specific variance observed between replicates from the same individual. To decompose these two variance components, we used a linear mixed effects model where we treated the individual as a random effect. As before, we utilized the inverse variance weights obtained from the voom approach and fitted the model using log-likelihood instead of a restricted maximum likelihood approach. We tested the null hypothesis that the variance of the random effect is zero. Rejection of the null hypothesis implied that there was significant inter-individual variance in the expression of the given transcript. We adopted a simulation-based approach using an exact likelihood ratio test implemented in RLRsim R package (Scheipl et al., 2008). Multiple-hypothesis correction was applied to RNA expression and ribosome occupancy p-values separately using Holm’s method.

## Cis-QTL identification

Association between gene expression and the genotype at each variant position located in the exons of the APPRIS transcripts was tested in the set of 21 unrelated Yoruban individuals using PLINK v1.07 (Purcell et al., 2007). For each transcript, replicate gene expression measurements were averaged for this analysis. The expression values were regressed on variant genotypes assuming an additive genetic model where genotype was coded as 0,1, or 2 copies of the alternate allele and restricting the testing to variants with a minor allele frequency >10% in the 21 unrelated Yoruban individuals.

## Genetic Determinants of Variability in Ribosome Occupancy: Defining upstream open reading frames

We used AUG and CUG as potential start codons, and UAG, UAA and UGA as potential stop codons. We decided to include non-canonical CUG starts based on two lines of evidence. First, CUG initiation has been reported in few well-documented cases such as FGF2, VEGF, c-myc and MHC class I transcripts (Hann et al., 1988; Meiron et al., 2001; Schwab et al., 2003; Vagner et al., 1996). Second, recent studies mapping genome-wide translation initiation sites have suggested that upstream translation initiates frequently from non-AUG codon, most prominently at CUG sites (Ingolia et al., 2011; Lee et al., 2012). Finally, we compared the uORFs in each individual against the uORFs found in the reference genome and labeled each uORF as gained or lost compared to the reference.

## Testing the effect of uORF events on ribosome occupancy

To group individuals by uORF differences on a given transcript, we first determined all possible combinations of uORF gain/loss events. For each set of uORF events, we assigned values of 0, 1, or 2 to each individual depending on how many alleles of that individual match the particular combination. We then tested whether the copy number of the uORF variants affects ribosome occupancy of the main coding region using two approaches. In the first approach, we used linear regression. In the second, more conservative approach, we fitted a linear mixed model assuming the difference in cell lines is an individual-specific random effect, i.e. treating the different cell lines of the same individual as “technical replicates”.

## The effect of Kozak region sequence on translation efficiency

We defined the Kozak region as the six nucleotides preceding the start codon and the two nucleotides following the start codon. We extracted the nucleotide sequence of this region from all annotated APPRIS transcripts and built a position weight matrix (PWM), which recapitulated the known Kozak sequence (Figure 5A). We tested whether the nucleotide content of the Kozak sequence affected translation efficiency using the Kruskal-Wallis test. Specifically, we tested whether transcripts split into four categories based on the nucleotide at a given position has the same translation efficiency. We corrected the p-value from this test using Bonferroni correction for the 8 tests (number of positions) that were performed.

## Association between Kozak region genetic variants and ribosome occupancy

Next, we collected all SNPs that intersect annotated Kozak regions. We scored the variant and the reference Kozak sequence using the PWM matrix obtained above. We coded each variant by the PWM score change and assumed an additive relationship between different positions in the Kozak region and copy number of the allele. We then tested whether the variants in the Kozak regions affect ribosome occupancy of the main coding region using a linear model (lm function in R; see discussion for uORFs for comparison to a linear mixed effect model). We utilized the inverse variance weights obtained by the voom method as before. For all Kozak variants affecting ribosome occupancy, we conducted the same association test using RNA expression level as the phenotype. As for the uORF analysis, we deemed RNA association to be not significant if the nominal p-value was greater than 0.05, or if the regression coefficient had the opposite sign.

## Luciferase reporter assays

To assay translation efficiency, we used of a bicistronic luciferase reporter construct (Jang et al., 2006). This construct has an SV40 promoter that drives the expression of a bicistronic transcript that includes both the firefly and renilla luciferase. While the renilla luciferase translation is cap-dependent, firefly luciferase has an Hepatitis C virus (HCV) internal ribosome entry site (IRES) that enables cap-independent translation.

Gene synthesis was carried out by GenScript and gene segments were cloned right in front of the start codon (ATG) of the renilla luciferase using the CloneEZ system (GenScript). A total 9 constructs were generated for four genes. The bicistronic constructs were transfected into HEK293 cells using lipofectamine 2000 (Invitrogen) in a 96 well format following manufacturer’s guidelines. Cap dependent translation was calculated by taking the ratio of renilla (cap) to firefly (HCV IRES) luciferase activity and derived from 5 replicate experiments. The HCV-IRES dependent translation of firefly luciferase accounted for differences in RNA expression and transfection efficiency. Outlier detection was carried out as described (Jacobs and Dinman, 2004). The differences between renilla to firefly luciferase ratios were assessed using Welch two sample two sided t-test.

## Data Access

The sequencing data reported in this paper has been submitted to NCBI GEO database. Accession number: GSE65912.

## Acknowledgements

CC is supported by Child Health Research Institute, Lucile Packard Foundation for Children’s Health and the Stanford CTSA grant number UL1TR000093. GWB is supported by a Benchmark Stanford Graduate Fellowship. SC is supported by a Stanford Transformative and Innovation grant. DS was supported by NIH/NHGRI T32 HG000044 and the Genentech Graduate Fellowship. Hua Tang is supported by the NIH grant GM07305907. This research was supported by NIH grants 1U01HG00761101, HG007611, and HG006996. We would like to thank Basar Cenik, Douglas H Phanstiel, Robert J Nichols for comments. We would like to thank Ghia Euskirchen for sequencing support. Illumina sequencing services were performed by the Stanford Center for Genomics and Personalized Medicine.

## Author contributions

CC, MPS designed the study. CC carried out ribosome profiling experiments and coordinated the computational analyses. GWB carried out uORF analysis, SC contributed to ribosome occupancy QTL analysis, BA developed visualization tools, HT contributed to RNA-Seq analysis. ESC performed luciferase reporter assays, FG generated RNA-Seq data, DS carried out cell culture/maintenance. ER helped optimize the ribosome profiling protocol, CA, and Hua Tang provided statistical analysis guidance. CC, ESC, CA, and MPS wrote the manuscript with input from all authors.

---

**Figure S1.**
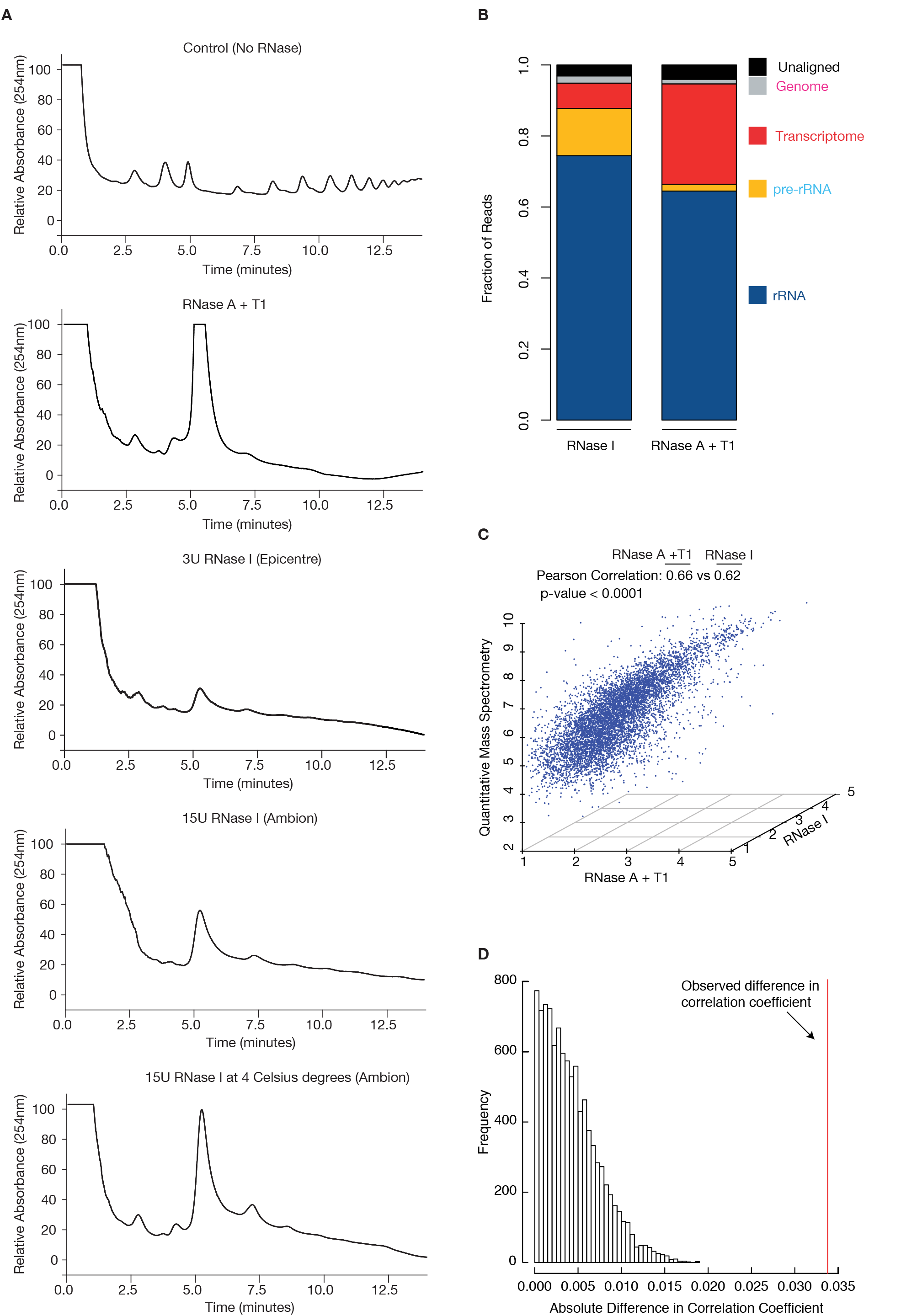

**Figure S2.**
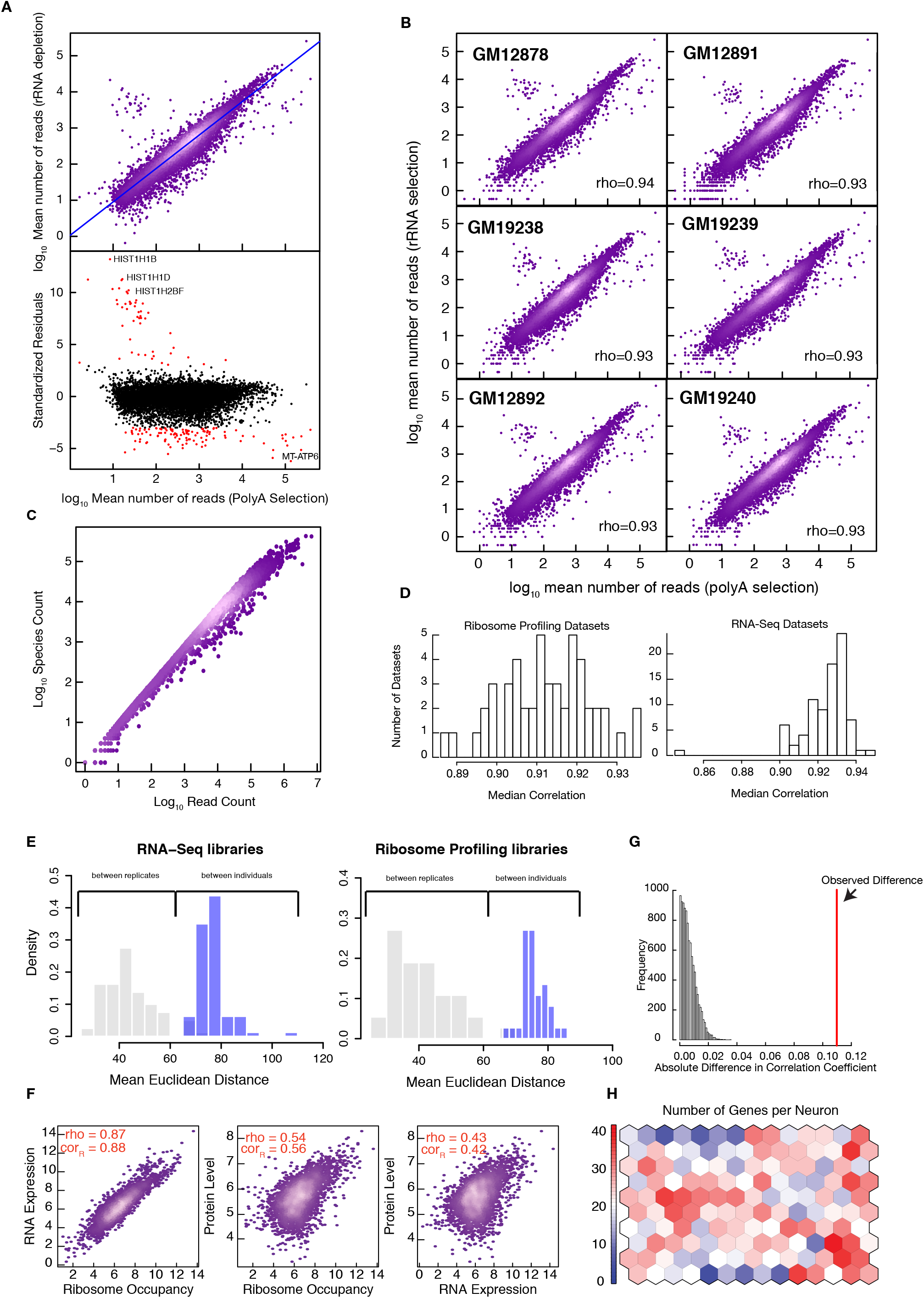

**Figure S3.**
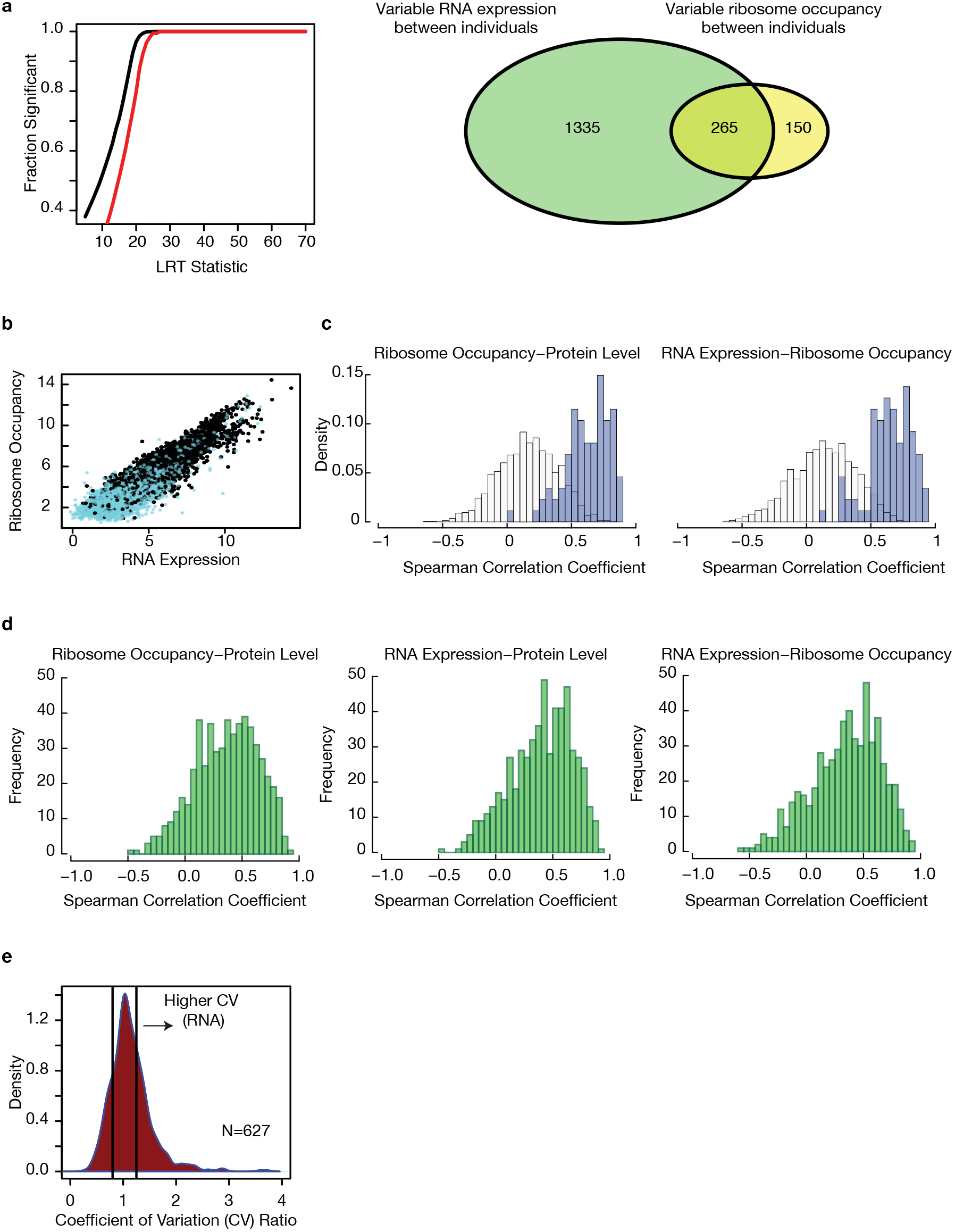

**Figure S4.**
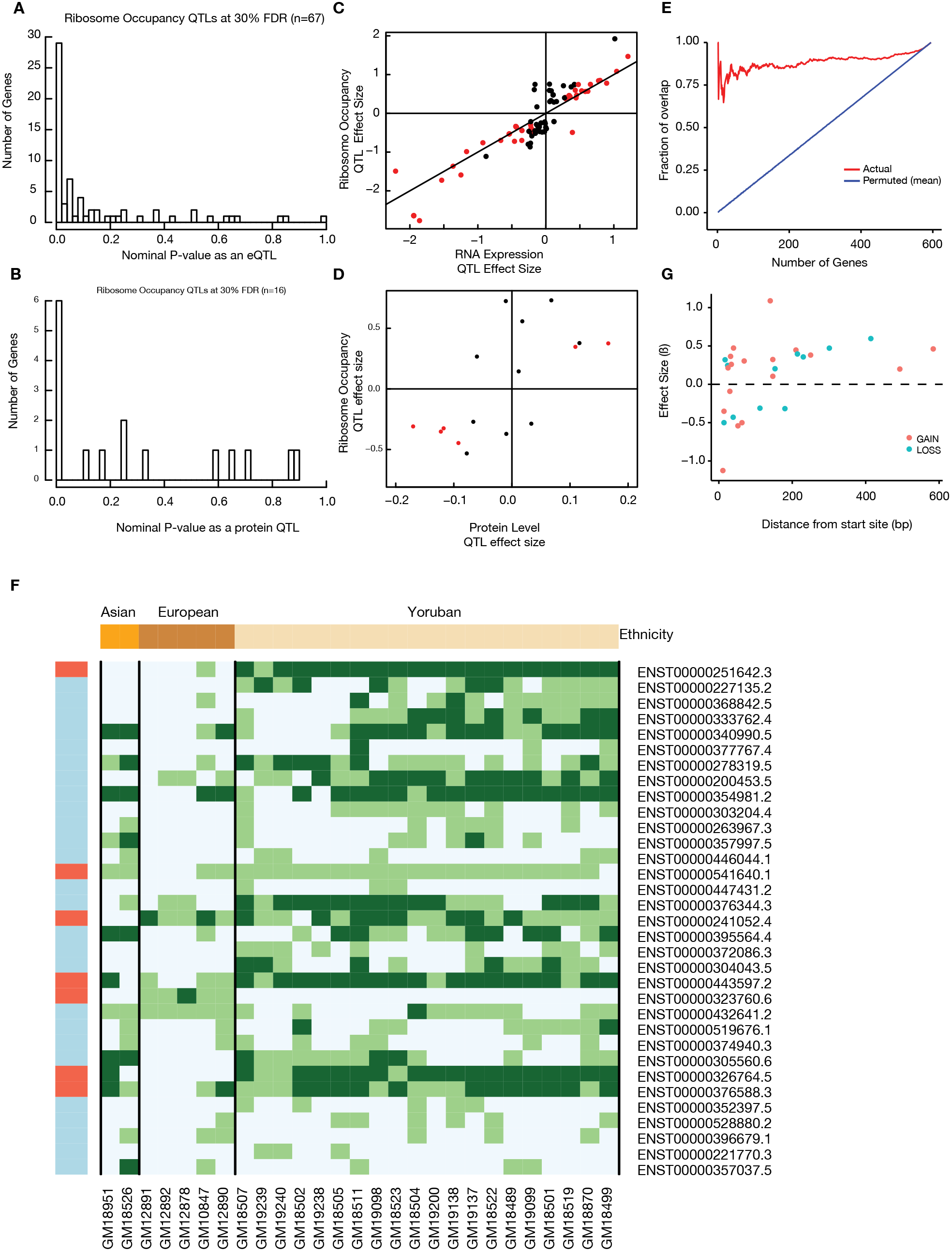

**Figure S5.**
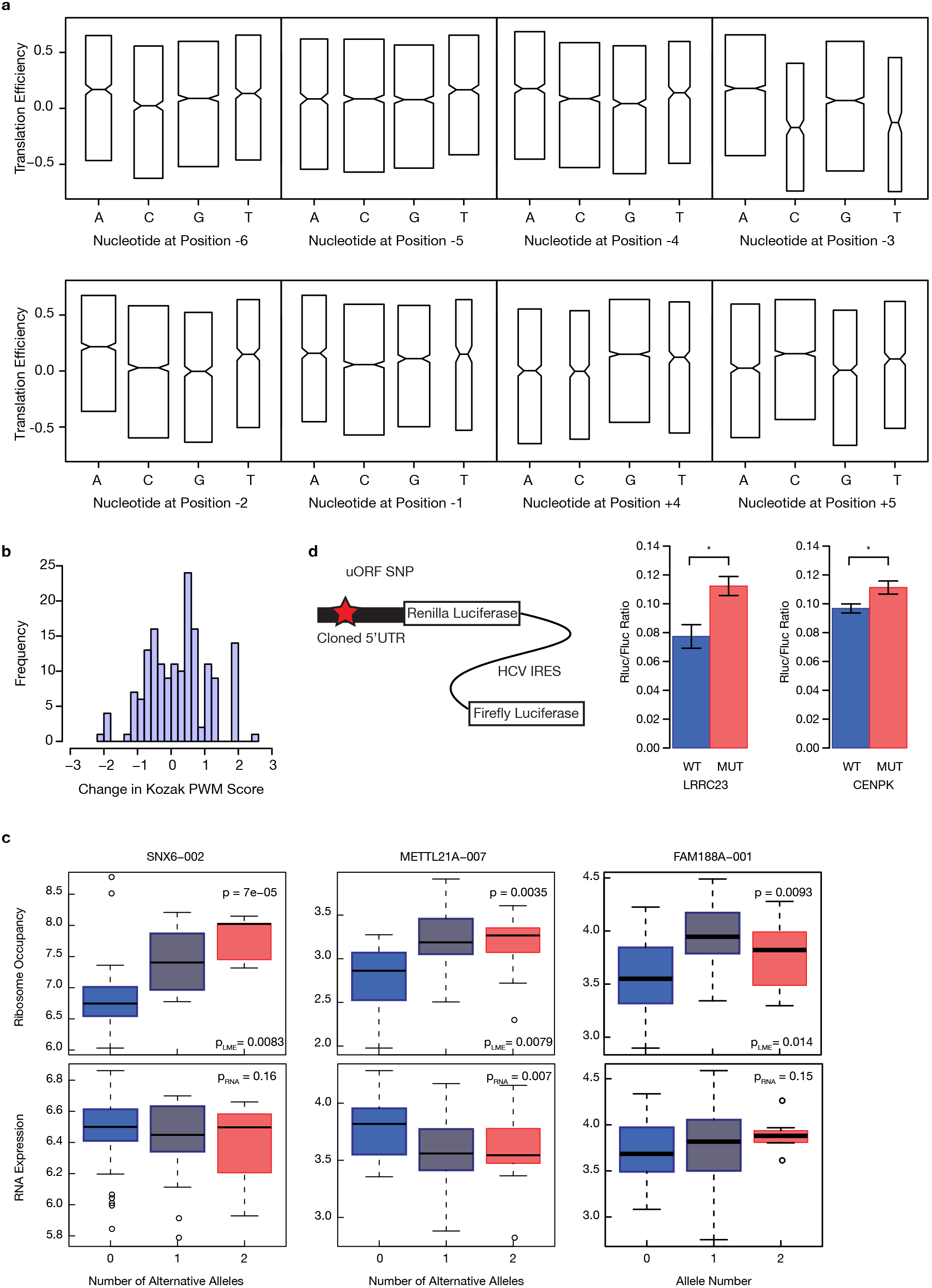

